# Global metabolome changes induced by environmentally relevant conditions in a marine-sourced *Penicillium restrictum*

**DOI:** 10.1101/2023.09.06.556477

**Authors:** Van-Tuyen Le, Samuel Bertrand, Marion Brandolini-Bunlon, Emmanuel Gentil, Thibaut Robiou du Pont, Vony Rabesaotra, Gaëtane Wielgosz-Collin, Aurélie Mossion, Olivier Grovel

## Abstract

Marine fungi have been found in all habitats and are able to adapt to their environmental niche conditions. In this study, a combination of LC-HRMS and GC-MS analytical approaches was used to analyse the whole metabolic changes of a marine sourced *Penicillium restrictum* strain isolated from a marine shellfish area. The *P. restrictum* MMS417 strain was grown on seven different media including an ecological one with two different water sources (synthetic sea water and distilled water) conditions following the OSMAC approach. Extracts of all media were analysed by LC-HRMS (lipids and specialised metabolites profiling) and GC-MS (fatty acids profiling). Aquired data were analysed using a multiblock strategy to highlight metabolic modification in regards to water conditions and to environmentally relevant conditions (mussel-based culture medium). This revealed that fatty acid composition of lipids was the most altered part of the explored metabolisms either looking to water effect and to environmentally relevant conditions. In particular, data showed that *P. restrictum* MMS417 is able to produce lipids that include fatty acids usually produced by the mussel itself. This study also provides insight into the *P. restrictum* adaptation to marine salinity through fatty acids alteration. and shows that lipid metabolisms if far more altered in an OSMAC approach than the specialized metabolism. This study finally highlights the need for using environnement-mimicking culture conditions to reveal the metabolic potentialities of marine microbes.

## 1. Introduction

Marine fungi have been found in all habitats and make a significant contribution to the marine environment [1]. They can be found as saprotrophs, parasites, or symbionts (epiphytic and endophytic) of marine macroorganisms, as a consequence of the evolution of fungal cell biology and feeding strategies [2, 3]. In the dynamic marine natural environments, fungi must be able to adapt to the change from the local environment to survive and colonize a niche. This could be considered as a physiological adaptation to their environment or as a response to abiotic stresses (rapid modification of environment) [4, 5]. Furthermore, in ecological niches fungi are also in competition or interactions with other micro- and/or macro-organisms, such as bacteria [6, 7], microalgae [8], seaweeds or macroalgae [9], sponges [10], or molluscs [11-13]. In a natural product perspective, biotic or abiotic stresses are responsible for activating specialised metabolite biosynthetic gene clusters (BGCs) yielding the production of stress-induced or activated specific compounds.

Considering marine-sourced fungi, salinity represents one of the main environmental factors to be considered to achieve one step further in understanding adaptation to marine environment. As one well documented effect in yeasts, salinity affects the membrane lipid composition of fungi, and consequently, its fluidity, in particular, through modifications in the type of fatty acyl chains, the sterol content, and the nature of the polar phospholipids head-groups [14-20]. However, up to now, no study trying to show an overall effect on the global metabolic content was initiated in fungi. A second factor to be considered for the understanding of the metabolome expression and regulation in marine fungi inhabiting holobionts is the influence of their micro-environment within their host. In this way, it has been previously shown that fungi cultured on host-derived media can produce specialised metabolites which were not observed in other conditions [21-24]. It can therefore be hypothesised that these compounds are involved in homeostasis or defence within their holobiont of origin.

However, studying these effects in fungi remains quite challenging [25]. The culture based strategies are usually the most employed for chemical diversity exploration, such as OSMAC (One-Strain-Many-Compounds) [26] or microbial co-cultivation [27]. These strategies allow to reduce the significant discrepancy between the actual number of chemically characterized compounds obtained from microorganisms and the number of bioinformatically characterised BGCs [28, 29]. The OSMAC approach is now largely used to explore the effect of various factors on fungal specialised metabolisms [28], such as temperature [30], carbon source [30-32] and culture medium [33].

Recent trends in metabolomics tend to perform a global analysis at multiple layers from core metabolome to lipids and specialised metabolome [34]. This reflects the ability that has emerged in recent decades to characterise the chemical composition of microbial extracts using different high resolution strategies, such as gas chromatography coupled to mass spectrometry (GC-MS), liquid chromatography coupled to mass spectrometry (LC-MS) and/or nuclear magnetic resonance (NMR) [35, 36]. It is also the consequence of the use of chemometrics pipelines that are able to correctly fuse complex data from various instrumentations using the so-called “statistical multiblock approach” [37-39]. Thus, in the goal of studying fungal metabolic adaptation to specific environmental conditions, it is now possible to acquire high resolution data using different methods and then to analyse them together to get a global overview [40, 41].

The present study proposed to explore the metabolic adaptation of the mussel-derived fungus *Penicillium restrictum* MMS417 isolated from mussel flesh within shellfish farming area [42] to both salinity and environnementaly relevant culture conditions. Such strategy, using host derived culture conditions, is yet poorly applied to microorganism’s growth for fungal metabolism exploration, as such microorganisms are nearly exclusively grown in standard laboratory conditions. This is expected to be a simple way of mimicking the original ecological niche, to describe *in situ* fungal metabolism behaviour. Thus, an OSMAC strategy was used to explore the metabolome expression in response to various culture conditions. In this way, six traditional culture media [33] were used along with a mussel flesh based medium [22]. This latter culture medium may be considered as an environmentally relevant - or ecological - medium [23, 24, 43]. In addition, each medium was studied as a marine-mimicking medium using synthetic sea water (SSW) or as an artificial laboratory medium using distilled water (DW). Thus, all together, the *P. restrictum* MMS417 metabolism corresponding to those 14 culture conditions was explored using specialised metabolitess [44] and lipid [31, 45] profiling by LC-HRMS and fatty acid lipid content profiling by GC-MS [46]. The multiblock analysis of the acquired data aimed to highlight specific metabolic changes in the *P. restrictum* MMS417 strain according to the use of SSW and of mussel extract in the culture medium.

## 2. Materials and methods

### 2.1. Fungal Material

The studied strain, *Penicillium restrictum* MMS417, was isolated from a blue mussel *Mytilus edulis* sampled in January 1997 at Port Giraud on the Loire estuary in France [42]. The strain is preserved at the laboratory ISOMer UR2160, Nantes Université, France. The strain has been identified after sequencing of *β*-tubulin gene and the internal transcribed spacer (ITS) and regions of the rDNA after nucleotide BLAST search (GenBank accession number for MMS417:KU720404, KU720398).

### 2.2. Culture Media Preparation

Fungal solid cultures were performed in Petri dishes (10 cm diameter) containing 15 mL of agar-based medium in 14 different media. Medium composition was as follow: DCA (Dextrose 40 g/L, enzymatic digest of Casein 10 g/L, Agar 15 g/L, Difco), MEA (Malt Extract 20 g/L, peptone 1 g/L, glucose 20 g/L, Agar 20 g/L), PDA (Potato extract 4 g/L, Dextrose 20 g/L, ZnSO_4_.7H_2_O 0.01 g/L, CuSO_4_.5H_2_O 0.005 g/L, Agar 15 g/L), YES (Yeast Extract 20 g/L, Sucrose 150 g/L, agar 20 g/L), MES (Mussel Extract 20 g/L, Sucrose 150 g/L, agar 20 g/L), CYA (Czapek concentrate 10 mL, Yeast extract 5 g/L, K_2_HPO_4_ 1 g/L, sucrose 30 g/L, Agar 15 g/L), Czapek concentrate (NaNO_3_ 3 g/L, KCl 0.5 g/L, MgSO_4_.7H_2_O 0.5 g/L, FeSO_4_.7H_2_O 0.01 g/L, ZnSO_4_.7H_2_O 0.01 g/L, CuSO_4_.5H_2_O 0.05 g/L), KMS (MgSO_4_.7H_2_O 2.4 g/L, NH_4_NO_3_ 2.4 g/L, Tris (tampon) 1.21 g/L, Agar 20 g/L). Details for the preparation of the mussel-derived medium (MES) are presented in a previous study [22]. Two osmotic conditions were prepared on the basis of the presence or absence of synthetic seawater (36 g/L of salinity – Coral Reef^®^ salts).

### 2.3. Fermentation and Extraction for OSMAC Approach

Fungal cultures were carried out in four replicates for each medium. Inoculation on Petri dishes was made from stock cultures of the strain stored at 20 °C and transplanted on MES medium 10 days before inoculation. Cultures were incubated at 27 °C for 10 days under natural light. Extraction of metabolites was performed on standardised plugs sampled in three distinct places of each culture (at the point of central impact, on the outskirts of the colony, in the periphery but in contact with an adjacent colony). Fungal biomass and agar layer were then extracted together to obtain both intra- and extracellular metabolites. Samples were gathered and extracted twice with 1.5 mL of CH_2_Cl_2_/EtOAc 1:1 (*v*/*v*) for each culture, sonicated for 30 min and filtered on regenerated cellulose filters 0.45 *µ*m (Sartorius) before drying. Media without fungi were also extracted following the same protocol and used as controls (blanks samples).

### 2.4. HPLC-MS analyses of specialised metabolites

HPLC-UV-(+/–)-HRESI-MS analyses were performed on a UFLC-MS (IT-TOF) Shimadzu instrument, using a Kinetex C18 column (2.6 *µ*m, 2.1 × 100 mm, Phenomenex) and following previously described conditions [44]. Samples (5 *µ*L) were injected at a concentration of 1 mg/mL in MeOH. A mixture of all extracts at the same concentration was also prepared as quality control (QC) sample. All samples were analysed randomly, and MeOH blanks and QC samples were regularly injected during the analysis sequence [47].

### 2.5. HPLC-MS analyses of lipids

HPLC-(+/–)-HRESI-MS analyses were also performed on a UFLC-MS (IT-TOF) Shimadzu instrument, using a Kinetex C18 column (2.6 *µ*m, 2.1 × 150 mm, Phenomenex). The conditions were previously described by [31]. Samples (2 *µ*L) at a concentration of 1 mg/mL in isopropanol were injected. As for the analyses of specialised metabolites, QC samples were prepared, and all samples were injected randomly, and isopropanol blanks and QC samples were regularly injected during the analysis sequence [47].

### 2.6. Processing of HPLC-MS data

The HPLC-(+/–)-HRESI-MS chromatogram raw data files were converted to *.netCDF files by using LCMS solution (version 3.60 – Shimadzu). Automatic peak picking on the *.netCDF files was achieved using MZmine 2 [48] with appropriate parameters (Table S1). All data files including media controls and MeOH blanks were examined to determine a minimum noise level threshold.

The results were exported as a *.csv file containing all peaks observed and referenced by their mass to charge ratio (*m/z*) and retention times (t_R_) together with their respective peak areas in each sample. The generated matrix was cleaned by removing all peaks abundantly found in blank samples (culture media extracts) and technical blank samples (MeOH injections).

### 2.7. Preparation of Fatty Acid Methyl Esters – FAMEs – and *N*-Acyl Pyrrolidides – NAP for GC-MS analyses

Fatty acid methyl esters (FAME) were prepared by submitting the crude extracts to transesterification during 5 h at 80 °C under reflux with methanolic hydrogen chloride 3 N/MeOH/CHCl_3_ (5:3:1 *v*/*v*/*v*). *N*-Acyl Pyrrolidides (NAP) were prepared by direct treatment of the FAME with pyrrolidine/acetic acid (5:1 *v/v*) for 60 min at 85 °C under reflux.

### 2.8 GC-MS analyses

FAMEs were analysed using a GC-MS instrument (Hewlett Packard HP 6890-GC System, Agilent Technologies, Santa Clara, CA, USA) linked to a mass detector (HP 6890-E.I. 70 eV) equipped with a SLB-5^TM^ column (60 m × 0.25 mm × 0.25 µm). The carrier gas was helium at a flow rate of 1 mL/min. The temperature of the injector and detector were respectively set at 250 °C and 280 °C. One microliter was injected in splitless mode. The column temperature was held at 170 °C for 4 min and programmed to 300 °C at 3 °C/min for FAME analyses and at 200 °C for 4 min and increased by 3 °C/min up to 310 °C for NAP ones. The solvent delay was 9 min.

### 2.9 Processing of GC-MS data

The GC-MS chromatogram raw data files were manually inspected to characterise all FA detected in the chemical profiles using GC-MS solution (Hewlett Packard 1989-1997, G1701BA version B.00.00). Complete identification of the peaks was performed by interpretation of the MS spectrum obtained from the NAP. In parallel, peak integration was performed on the FAME profiles manually and aligned based in former identification to provide the GC-MS data matrix. The generated matrix was cleaned by removing all peaks abundantly found in blank samples (culture media extracts) and technical blank samples.

### 2.10 Statistical Analysis

All data analyses were performed using R 4.0.0 (CRAN). Data from HPLC-HRESI-MS profilings were subjected to QC correction using the statTarget package [49] using the following parameters: Frule 0, QCspan 0.75, degree 2, imputeM “KNN” and coCV 100. The corrected peak areas were further corrected to reflect the absolute compound amount in the fungus by multiplying it by the corresponding extract weight. A similar correction was applied to GC-MS peak area. All data analyses were performed using Unit-Variance (UV) scaling. The PCA was performed using the ROPLS package [50]. The ComDim data analysis [51] was performed using the R function “accps()” available from E. Tchandao Mangamana [52]. The MBPLS-DA was achieved using the packMBPLSDA package [38] and the variables were selected based on VIPc (cumulated variable importance in the projection) values.

### 2.11 Annotation of LC-HRMS data

The selected features based on VIPc were putatively identified taking into acount their accurate molecular masses. In details, measured accurate masses were used for compounds molecular formula determination considering a 6 ppm error, which was compared with the information available on the dictionary of natural products (DNP) database. UV spectra were used to suggest secondary metabolite classes.

## 3. Results

To assess the impact of the modification of culture conditions on the metabolism of *P. restrictum*, 14 culture media were used in an OSMAC experiment. The fungus was grown for 10 days using seven solid culture media (CYA, DCA, PDA, YES, MEA, KMS and MES), supplemented or not with synthetic seawater, *prior* to extraction. Tracking a global effect of the replacement of distilled water (DW) by synthetic seawater (SSW) needs to analyse all metabolic changes in the culture conditions, having in mind to get as much information as necessary on the fungal metabolism. Thus, after extraction of the culture medium and the fungal biomass together, all extracts (n = 56) were profiled focusing on specialised metabolites (SM), lipids, and fatty acids (FA) (Figure 1). Two different LC-HRMS protocols were used for first profiling specialised metabolism [44] and then lipids [31]. In parallel, an aliquot of each extract was subjected to transesterification to obtain the full FA composition of lipids.

**Figure 1.**
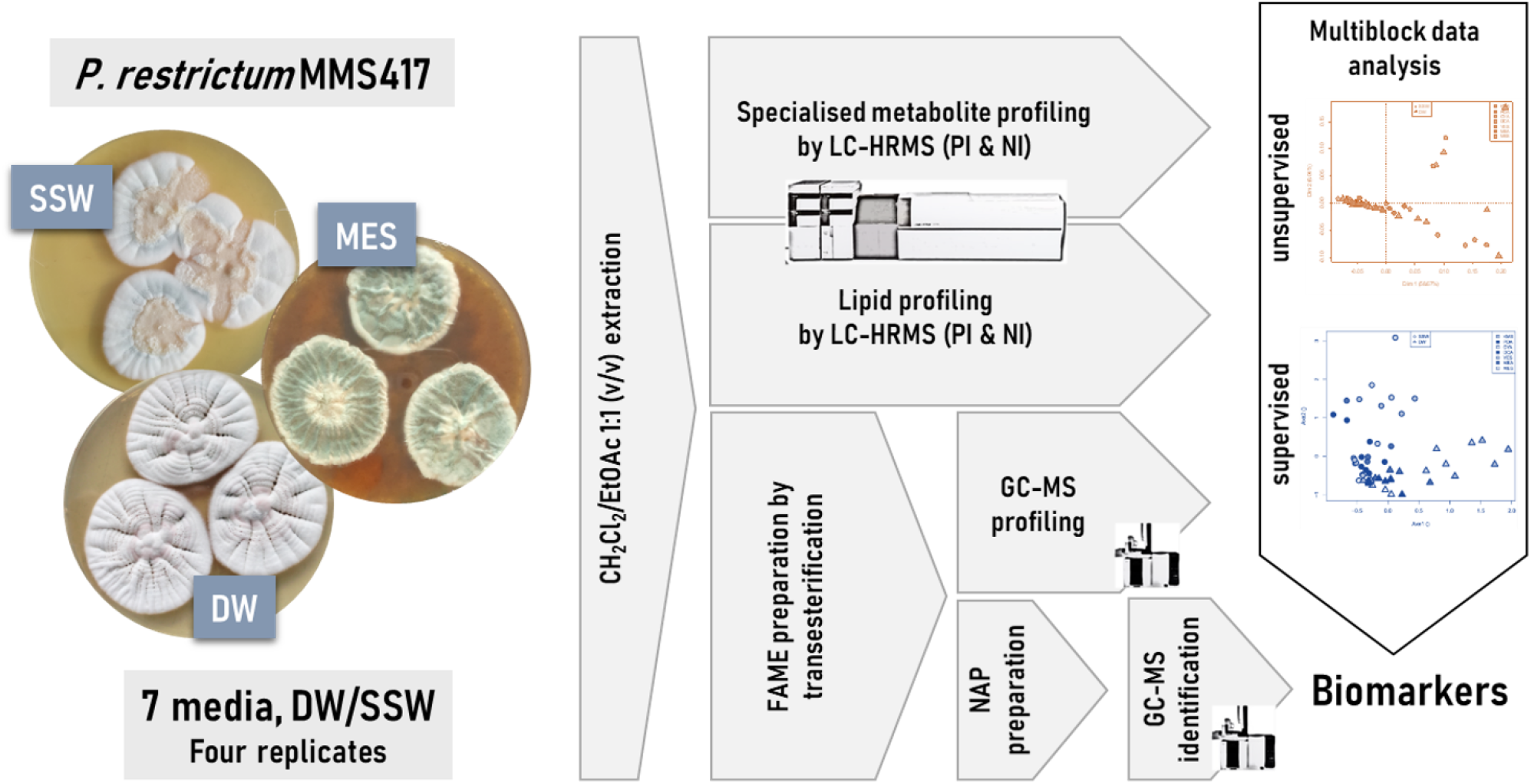
Experimental design used to assess the effect of seawater and environnementaly relevant culture conditions on the metabolome and lipidome of *Penicillium restrictum MMS417*. SSW: synthetic sea water, DW: distilled water, MES: mussel-derived culture medium, PI: positive ionisation, NI: negative ionisation, NAP: *N*-Acyl Pyrrolidides.

### 3.1 General profiling

As a first step, a simple LC- and GC-MS-based metabolomics analysis was performed to compare the metabolic profiles of all culture conditions. Figure 2 provides an overview of all acquired metabolic profiles (LC-MS and GC-MS). This gave a snapshot of the chemical composition of the strain when grown on different media, from which it can be easily visualised that the chromatogram profiles showed variations according to culture conditions.

**Figure 2.**
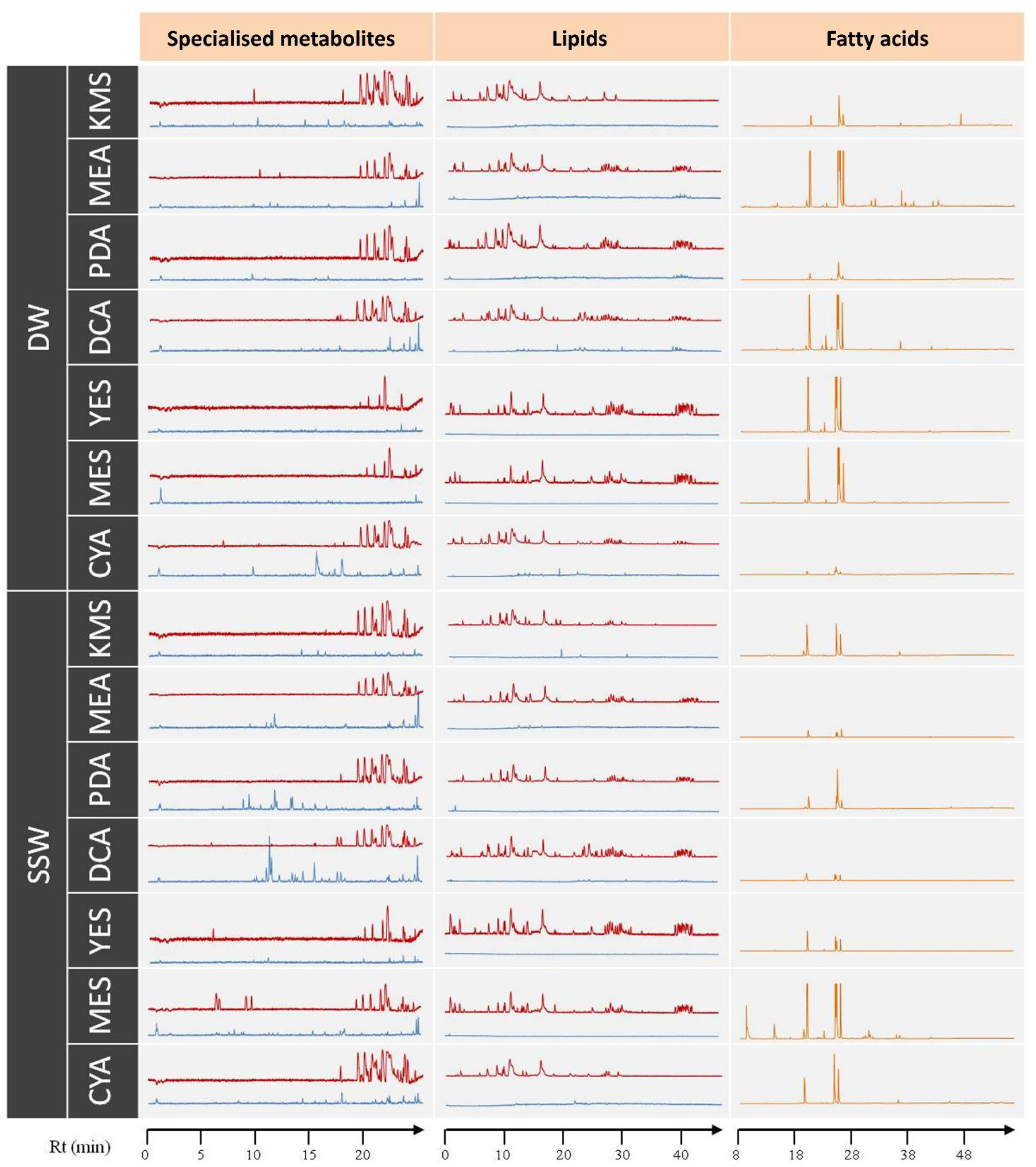
LC-HRMS (base peak chromatograms) and GC-MS (total ionic current chromatograms) profiles of the extract from *Penicillium restrictum MMS417* among all 14 cultures conditions: CYA, DCA, PDA, YES, MEA, KMS, and MES using either DW or SSW. The red line corresponds to the positive ionisation mode and blue line to the negative mode.

As a general observation on the specialised metabolite profiles in the positive ionisation mode, the majority of the peaks appeared at the end of the chromatograms (18-25 min). Interestingly, there were some particular peaks observed mostly in the profiles corresponding to MES-SSW extracts in comparison to the other media (5-10 min). In the negative ionisation mode, many peaks appeared only in the SSW profiles in comparison to those from the distilled water cultures (except CYA-DW). This was particularly observed in the case of PDA-SSW and DCA-SSW extracts.

A similar observation was achieved on lipidome. In the lipid profiles (Figure 2), the most polar lipids (*i.e*. FA) along with specialised metabolites eluted at the beginning of the chromatogram (approx. 2-18 min), phospholipids (PL) and glycolipids (GL) between 18 and 35 min, and finally neutral lipids (NL) (mostly triglycerides) between 38 and 45 min. Profiles obtained from both KMS and CYA media appeared clearly different with almost no PL, GL nor NL observed. Contrarily, all lipid categories were observed in profiles from MES and YES media. In addition, to these general lipidome composition observations, fatty acid compositions of lipids were also characterized leading to the observation that lipid composition was stable qualitatively except for GC-MS profiles obtained from MEA-DW and MES-SSW extracts wich showed a strongly different composition in comparison to others. In this way, *P. restrictum* MMS417 contained FAs of carbon chain lengths ranging from C12 to C26. The most common and abundant FAs were 16:0, 18:0, 18:1, and 18:2 which, taken altogether, often represent more than 90% of the total FA content of fungi [53-55]. These four FAs were present in all extracts in this study. Minor FAs (12:0, 14:0, 15:0, 16:1, 17:0, 17:1, 19:1, 19:0, etc.) were also detected, but represented less than 10% of the total FA content. However, the relative concentration of each individual fatty acid ranged from less than 1% of the total acid fatty acid content to over 47%, showing a high variability depending on the culture medium.

To further analyse the profiling data, multivariate unsupervised data analyses were performed using principal component analyses (PCA) on the following data from all samples: LC-HRMS profiles in positive ionisation mode (883 features), LC-HRMS metabolome profiles in negative ionisation mode (544 features), LC-HRMS lipidome profiles in positive ionisation mode (1748 features), LC-HRMS lipidome profiles in negative ionisation mode (10535 features), and GC-MS FA composition (34 features). In all PCAs (Figure 3), the first axe represented 24% (LC-HRMS metabolome profiles in positive ionisation) to 61% (LC-HRMS lipidome profiles in negative ionisation) of the variability, and a moderate clustering according to the culture media could be observed using the first two components (too many culture conditions were explored to expect visible clustering). All specialised metabolome and lipidome analyses revealed some discrimination of the MES and YES extracts (independently of the presence of synthetic sea water), and FA profiling also highlighted clear differences for these two culture conditions. In addition, the second axe of the lipidome data in positive ionisation highlighted a KMS culture medium specificity.

**Figure 3.**
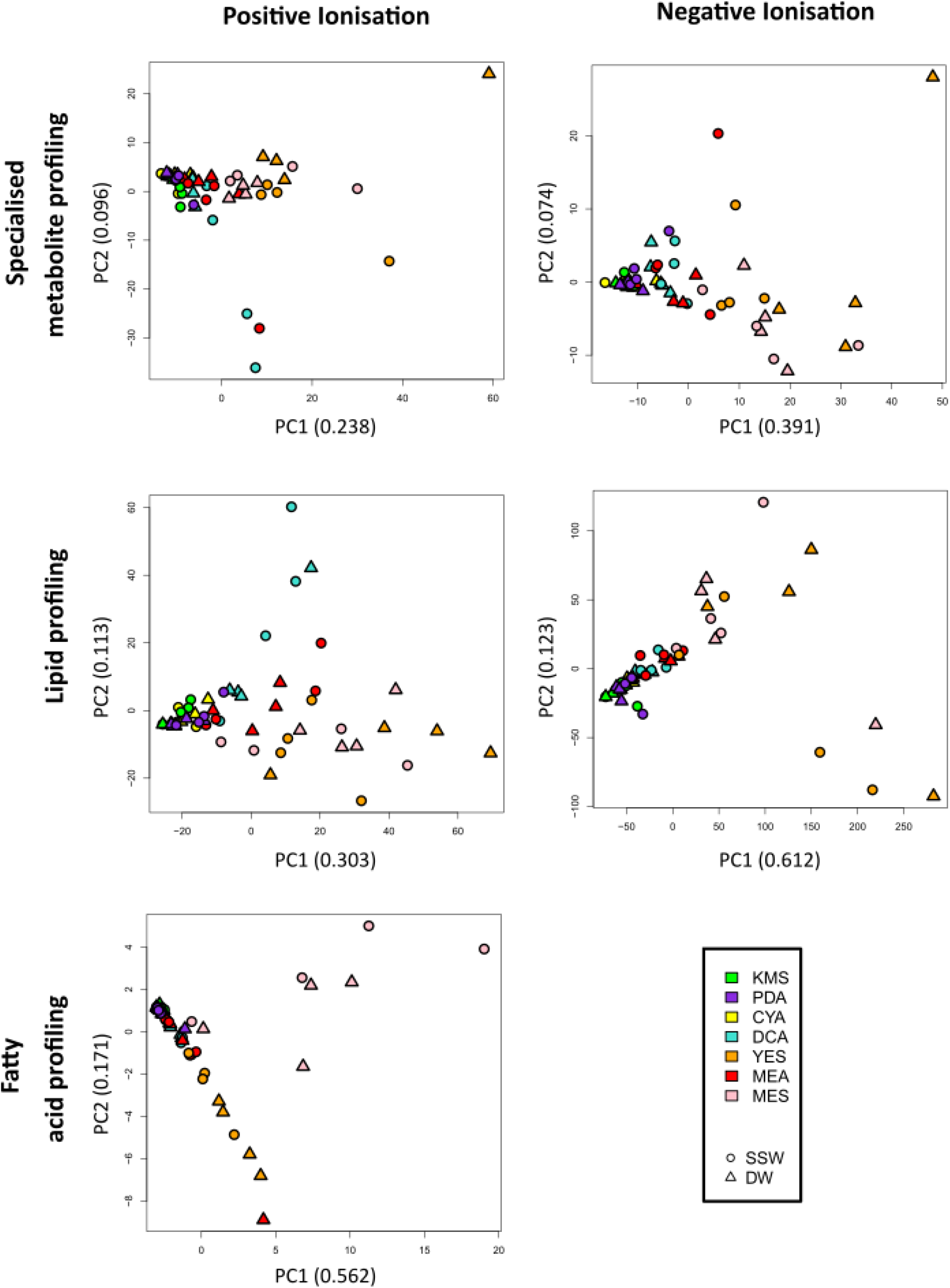
PCA score plot (UV scaling) of all acquired data (LC-HRMS and GC-MS) on extracts obtained from the 14 culture media (4 replicates). Triangles and circles correspond to DW and SSW, respectively.

However, the ability to conclude on a global culture medium effect using such a high amount of data was impaired by the difficulties to use at the same level the different data acquired on the extracts. Therefore, it could be interesting to analyse all data simultaneously. In such cases, a so-called multiblock data analysis can be performed [34].

Multiblock data analyses correspond to strategies allowing the co-analysis of different datasets acquired from the same samples, one block corresponding to one analytical method in our study. One major difficulty that multiblock strategies encounter is the block size differences effect. In our study, five blocks of data were acquired, having a high variation in data size. To overcome the block size differences effect, the unsupervised multivariate ComDim strategy (also called Common Components and Specific Weights Analysis – CCSWA) [39] was used to analyse the data. This multiblock strategy is still very rarely used to explore metabolomic data, particularly in the natural products field [41]. However, in addition to perform a simultaneous analysis of all data, it provides also informations about the importance of each data block.

The ComDim score plot (Figure 4AB) allowed visualising in an unsupervised way all the data self-structure. As observed in the individual data set (Figure 3), the MES and YES culture media (using both DW and SSW) corresponded to the first dimension which accounts for 59% of the total variance of the data. This first dimension (Dim1) was mostly derived from the lipidome in the negative ionisation data block as observed in Figure 4C. The second dimension (Dim2), which accounted for 8% of the total variance, highlighted the MES specific effect. Interestingly, in addition to lipidome specificity, FA composition exhibited a high specificity in the MES medium. Exploring further the dimension of the ComDim analysis (Figure 4B) highlighted the tendency for culture media to cluster. However, no SSW cluster were observed, suggesting the lack of a global salinity effect.

**Figure 4.**
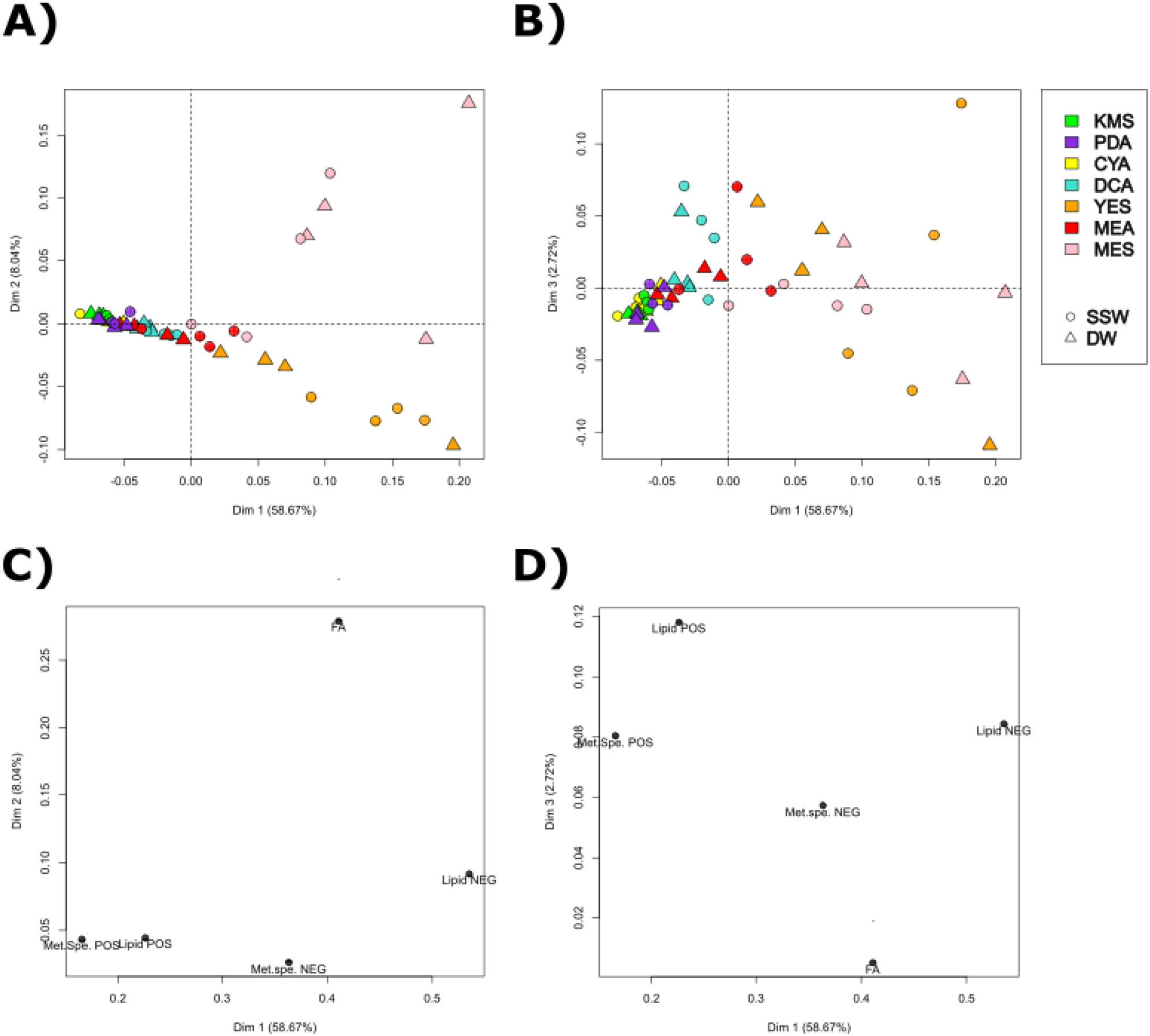
Multiblock unsupervised data analysis using ComDim (extract weight correction and UV scaling) of all acquired data (LC-HRMS and GC-MS). A) Score plots of dimensions 1 and 2 and B) Score plots of dimensions 1 and 3; C) Block influence on the separation (salience on the separation) of dimension 1 and 2 and D) Block influence on the separation (salience on the separation) of dimensions 1 and 3.

### 3.2 Effect of synthetic seawater on the metabolome expression

To investigate more in depth the effect of seawater on fungal metabolism (*i*.*e*. SSW ewtracts *vs* DW extracts), data were analysed in a supervised way using MBPLS-DA. Optimization of the number of PLS components indicated that only the first 2 should be kept to avoid overfitting [38]. The corresponding cross validated error rate of classification was 1.8 % ± 1.8 %. The resulting score plot (Figure 5A) showed a significant separation between medium based on DW and SSW. Exploration of the importance of each block in the separation (Figure 5B) indicated that all blocks impacted similarly the separation, however the most important block in the separation remained the lipid profiling acquired in negative ionisation and the less important block was the FAs composition.

**Figure 5.**
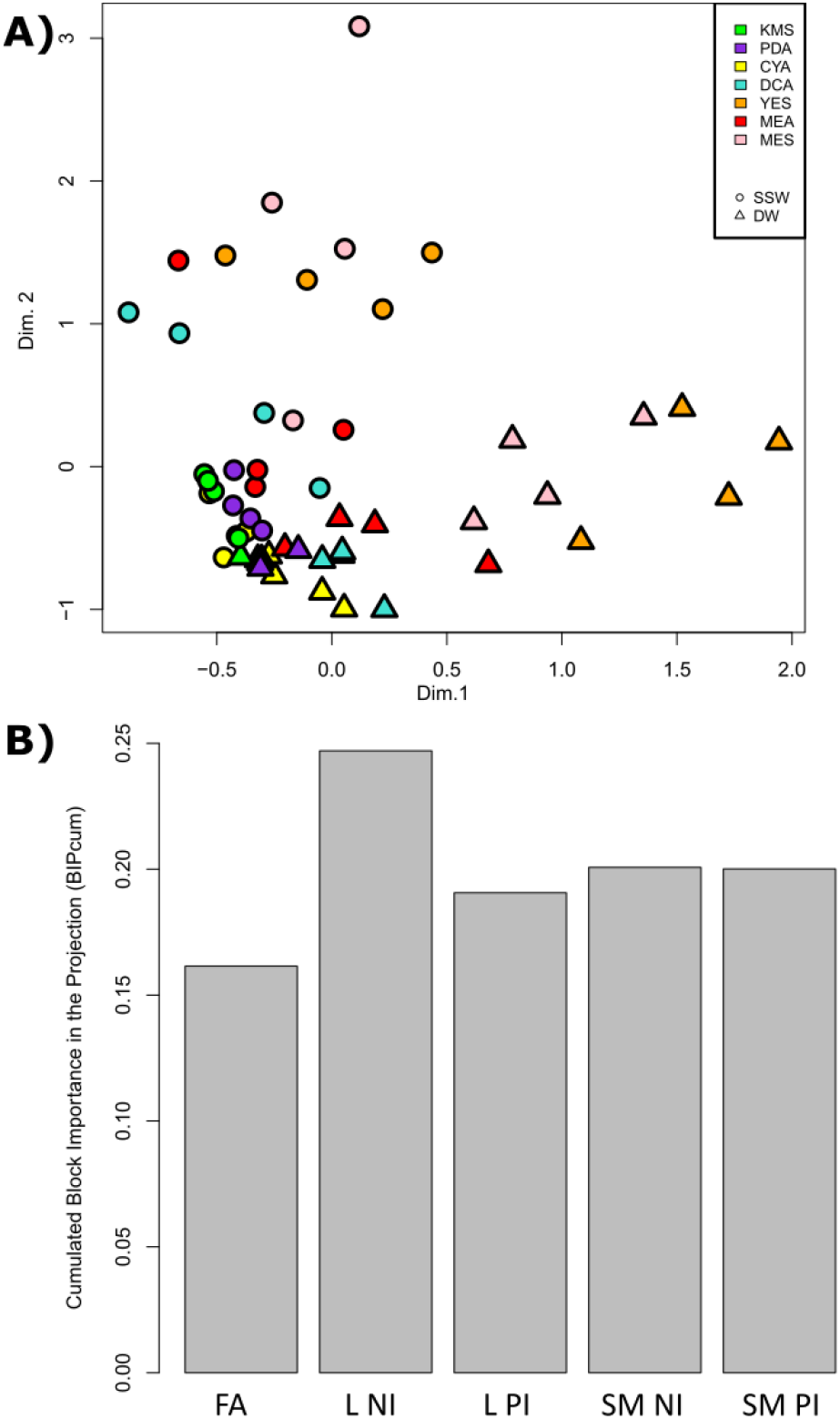
Multiblock supervised data analysis using MBPLS-DA (extract weight correction and UV scaling) of all acquired data (LC-HRMS and GC-MS) to discriminaite DW and SSW. A) Score Plot; B) Block importances in the projection. FA: fatty acid profiles by GC-MS; LP: Lipid profiles either in the negative (NI) or positive (PI) ionisation; SM: specialised metabolites profiles either in the negative (NI) or positive (PI) ionisation.

Based on VIPc values, features of interest were selected (Table 1). Among the Top30 features selected, besides fatty acids 16 features corresponded to ions detected by specialised metabolites profiling. The feature corresponding to *m/z* 427.3208 [M+H]^+^ eluted at 21.88 min was annotated as dankasterone B (C_28_H_42_O_3_, calculated *m/z* 427.3207, Δppm 0.23), a fungal steroid [56-58]. The feature *m/z* 482.2927 [M+H]^+^ eluted at 21.68 min was annotated as isomers of cytochalasin B_5_ (C_29_H_39_NO_5,_ calculated *m/z* 482.2901, Δppm 5.39), an hybride PKS-NRPS macrolide previously isolated from a marine-derived fungus *Phoma* sp. [59]. Some unidentified metabolites at *m/z* 209.0781 [M+H]^+^, 524.3151 [M+H]^+^, 363.0273 [M-H]^-^, 214.1452 [M-H]^-^, 429.287 [M-H]^-^, 437.2544 [M-H]^-^, 386.2595 [M-H]^-^, 484.3054 [M+H]^+^, 232.6385 [M+H]^+^ and two isomers at *m/z* 744.5735 [M+H]^+^, 744.5733 [M+H]^+^ did not find any hit from fungal natural products databases.

**Table 1.**
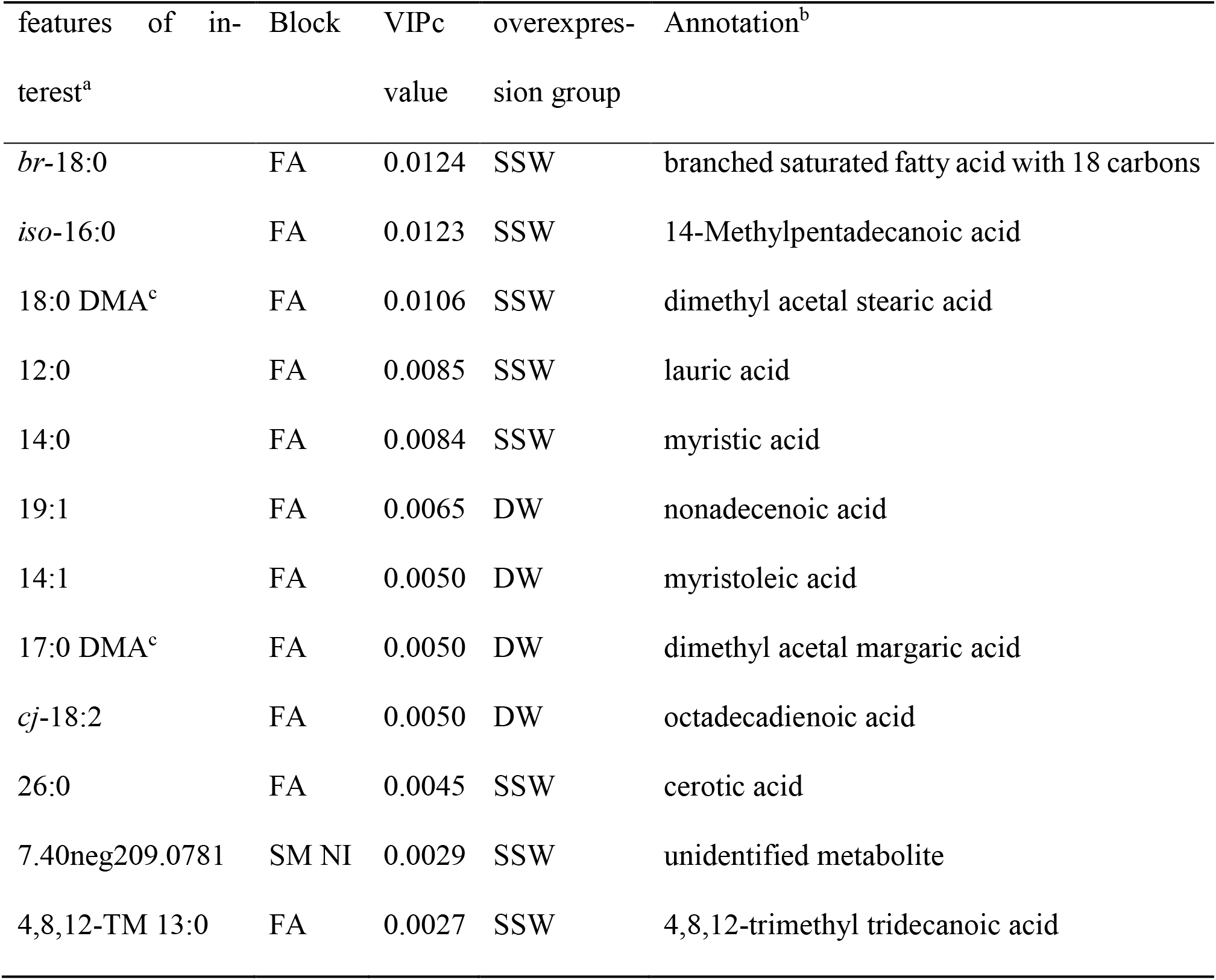
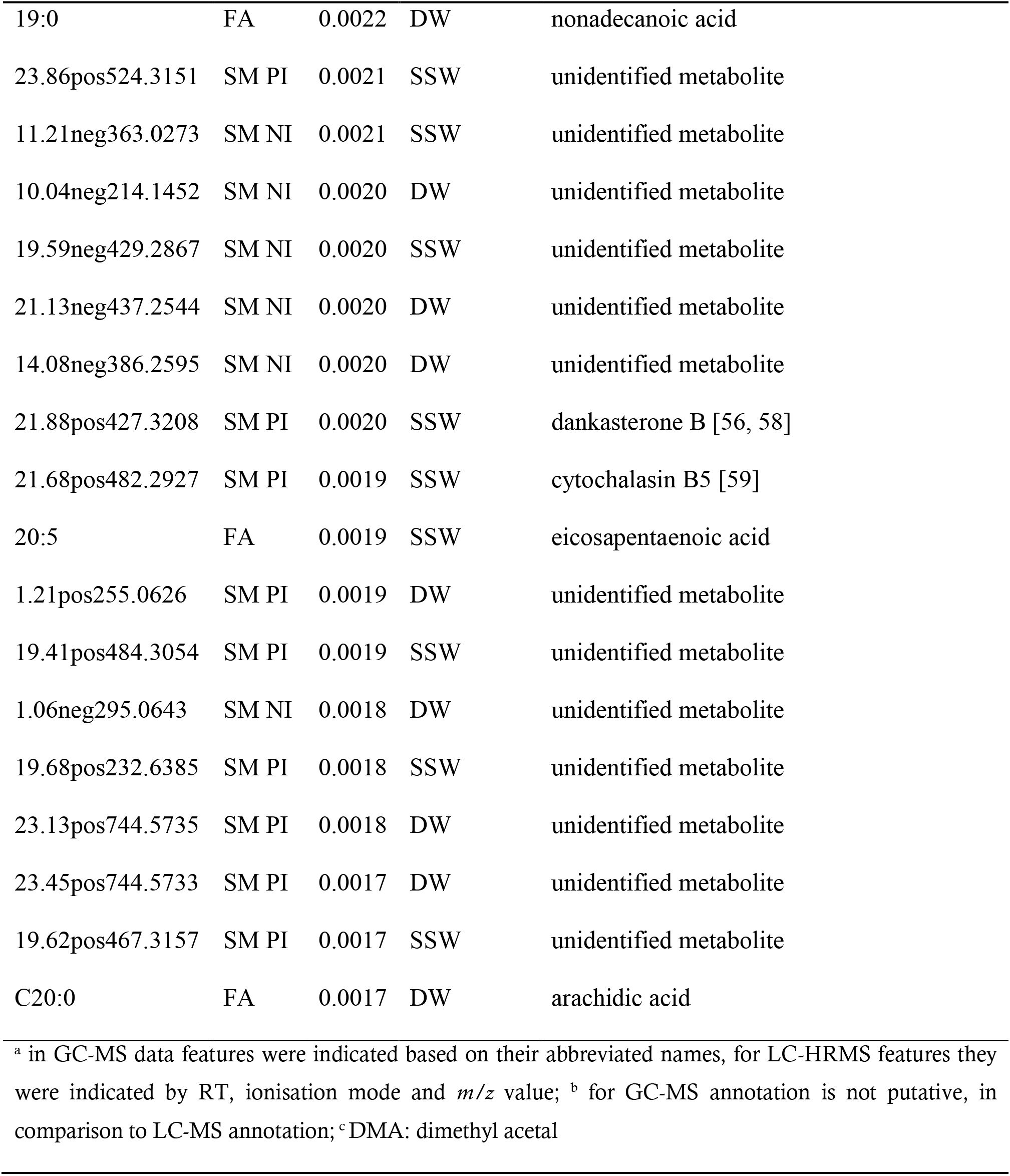
Most altered features (Top30) regarding DW and SSW differences highlighted by MBPLS-DA (Figure 6) and selected based on cumulated variable importance in the projection (VIPc).

The top10 VIPc variables related to this discrimination were FA. even if the FA block importance in the MBPLS-DA model was the smallest. This reflects that a large number of lipids observed in negative mode ionisation profiles are altered slightly but significantly. Concerning lipids, it was observed that their FA composition in the extracts using SSW culture media corresponded mostly to saturated fatty acids (SFA) (12:0, 14:0, 19:0, 20:0, 26:0), PUFA (*cj*-18:2, 20:5) and BCFA (*br*-18:0, 18:0-DMA, *iso*-15:0, 4,8,12-TM 13:0), whereas the one in the extracts from DW cultures corresponds to mono-unsaturated fatty acids (MUFAs) (14:1, 19:1), BCFA (17:0-DMA), PUFA (*cj*-18:2).

### 3.3. Effect of mussel components on the metabolome expression

As clearly observed in Figure 4A, the composition of *P. restrictum* MMS417 extract when grown on MES and YES medium exhibited specific chemical patterns without considering DW *vs* SSW effect. To observe such an effect on overall fungal metabolism, data were analysed in a supervised way using MBPLS-DA. The analysis was focused on highlighting differences between the MES culture medium in comparison to all other media, disregarding DW and SSW. Optimisation of the number of PLS components indicated that only the first 2 should be kept, allowing to obtain a classification error rate of 5.5 6% ± 5.56 %, and to avoid overfitting [38]. The resulting score plot (Figure 6A) showed a significant separation between the MES media and the others. Exploration of the importance of each block in the separation (Figure 6B) indicated that the FA block was the most important one, all other blocks being of similar importance.

**Figure 6.**
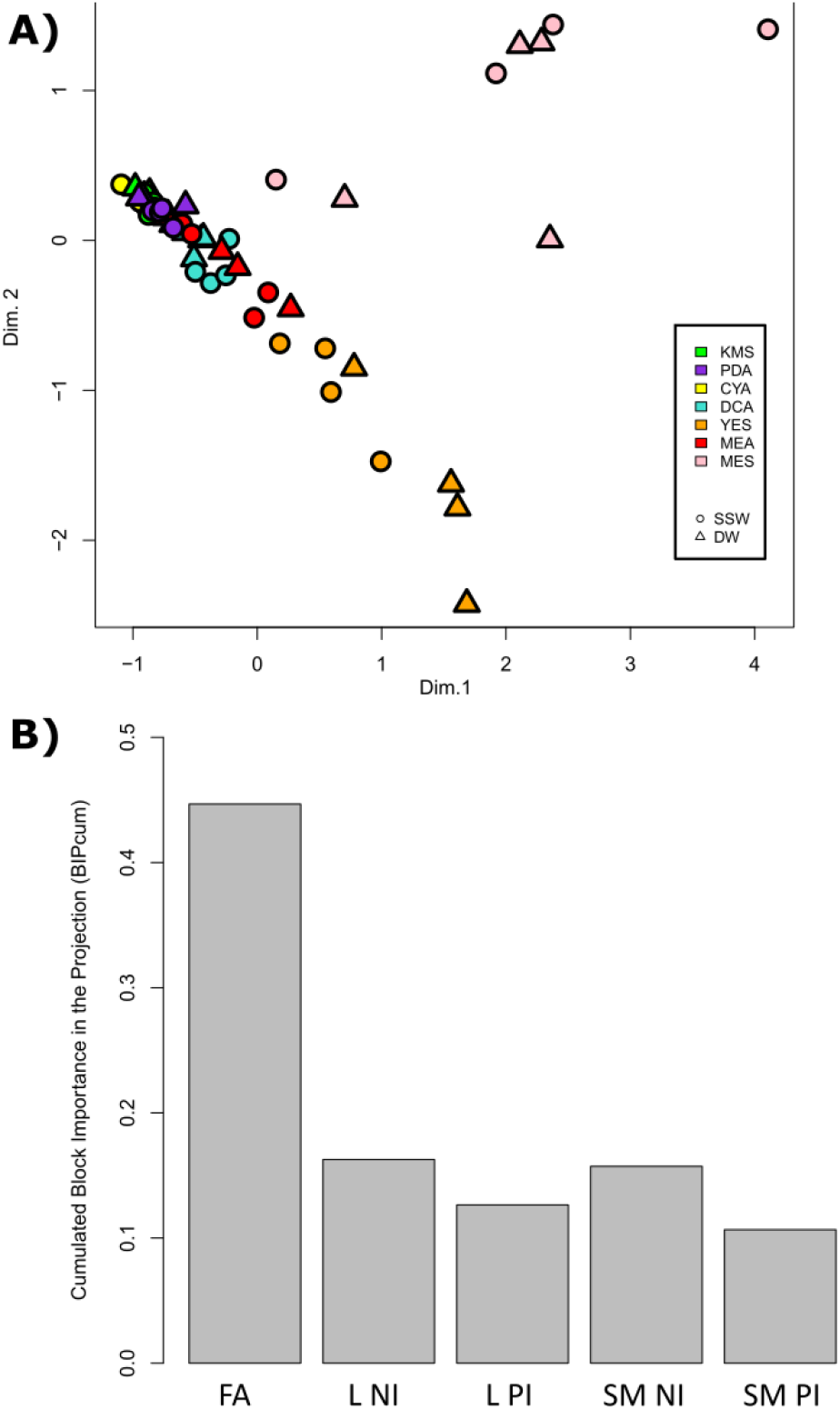
Multiblock supervised data analysis using MBPLS-DA (extract weight correction and UV scaling) of all acquired data (LC-HRMS and GC-MS) to discriminate the MES media from the other culture conditions. A) Score Plot; B) Block importances in the projection. FA: fatty acid profiles by GC-MS; L: Lipid profiles either in the negative (NI) or positive (PI) ionisation; SM: specialised metabolites profiles either in the negative (NI) or positive (PI) ionisation.

Based on the cumulated variable importance in the projection (VIPc), features of interest were selected (Table 2). Interestingly, the Top30 VIPc variables related to this discrimination were from the FA profiles except one from specialized metabolism. The only feature from another data block corresponded to the feature *m/z* = 415.2713 [M-H]^-^ eluted at 15.03 min and annotated as the tricarboxylic fatty acid – agaric acid (C_22_H_40_O_7_, calculated *m/z* 415.2697, Δppm 3.85). Interestingly, some of the FAs only present in the extracts obtained using the MES media were branched FAs (BCFA) or polyunsaturated FA (PUFA) such as 4,8,12-TM 13:0, 18:0-iso, 18:0-anteiso, 18:0-DMA, br-18:0, 20:4, 20:5.

**Table 2.** Most altered features (Top30) regarding MES media and all other culture conditions highlighted by MBPLS-DA (Figure 6) and selected based on cumulated variable importance in the projection (VIPc).

| Features<sup>a</sup> | Block | VIPc | Overexpressed in<br>MES | Annotation<sup>b</sup> |
| --- | --- | --- | --- | --- |
| **22.2** | FA | 0.0504 | yes | 8,13-docosenoic acid |
| **4,8,12-TM 13:0** | FA | 0.0494 | yes | 4,8,12-trimethyl-<br>tridecylic acid |
| **17.0.iso** | FA | 0.0493 | yes | isomargaric acid |
| **20.1** | FA | 0.0491 | yes | eicosenoic acid |
| **17.0.anteiso** | FA | 0.0442 | yes | anteisomargaric acid |
| **20.2** | FA | 0.0439 | yes | 6,11-eicosenoic acid |
| **20.4** | FA | 0.0439 | yes | eicosatetraenoic acid |
| **16.1** | FA | 0.0423 | yes | palmitoleic acid |
| **20.5** | FA | 0.0403 | yes | eicosapentaenoic |
| **16.0.iso** | FA | 0.0304 | yes | isopalmitic acid |
| **18.0.iso** | FA | 0.0287 | yes | isostearic acid |
| **15.0** | FA | 0.0284 | yes | pentadecylic acid |
| **18.0.anteiso** | FA | 0.0279 | yes | anteisostearic acid |
| **18.0.DMA<sup>c</sup>** | FA | 0.0268 | yes | dimethylacetal stearic<br>acid |
| **18.0.br** | FA | 0.0257 | yes | branched octadecanoic<br>acid |
| **16.0** | FA | 0.0218 | yes | palmitic acid |
| **18.0** | FA | 0.0170 | - | stearic acid |
| **14.0** | FA | 0.0142 | yes | myristic acid |
| **19.0** | FA | 0.0141 | yes | nonadecylic acid |
| **20.0** | FA | 0.0139 | - | arachidic acid |
| **17.0** | FA | 0.0108 | - | margaric acid |
| **18.1** | FA | 0.0106 | - | oleic acid |
| **22.0** | FA | 0.0088 | - | behenic acid |
| **12.0** | FA | 0.0087 | yes | lauric acid |
| **24.0** | FA | 0.0084 | - | lignoceric acid |
| **18.2** | FA | 0.0082 | - | linoleic acid |
| **17.1** | FA | 0.0043 | - | 9-heptadecenoic acid |
| **22.1** | FA | 0.0033 | yes | docosenoic acid |
| **19.1** | FA | 0.0031 | - | nonadecenoic acid |
| **15.03neg415.2713** | SM NI | 0.0012 | yes | agaric acid [60, 61] |

<sup>a</sup> in GC-MS data features were indicated based on their abbreviated names, for LC-HRMS features they were indicated by RT, ionisation mode and $m/z$ value; <sup>b</sup> for GC-MS annotation is not putative, in comparison to LC-MS annotation; <sup>c</sup> DMA: dimethyl acetal

## 4 Discussion

Several studies have previously demonstrated the high capacity of fungi to adapt their metabolic processes to environmental variables such as temperature and salinity conditions [16, 62-65]. It was proposed that fungi are able to reshape the molecular structure of their cell membrane through modification of their lipid composition such as modification of saturated, unsaturated, branched, or cyclic fatty acids contents in individual phospholipids [66, 67]. In addition, it is a now well-known fact that expression of biosynthetic gene clusters responsible for the production of specialised metabolites is under the control of various biotic or abiotic factors [28, 68]. In this way, many natural products have been discovered following culture screenings using the so-called OSMAC approach [26, 28]. In the present study, we envisaged to explore the global metabolome variations induced by media modifications by analysing both the lipidome including the FA composition of lipids, and the specialised metabolome of *P. restrictum*. Unlike many studies, the choice of growing conditions was dictated by ecological considerations to understand the influence of marine salinity and the microenvironment of the original holobiont of the fungal strain studied. This was achieved by using a medium based on mussel flesh and reconstituted with seawater, a medium that tended to come as close as possible - under laboratory conditions - to the original conditions in which the strain was naturally growing.

The supervised multiblock data mining strategy revealed a clear effect of addition of SSW in the culture medium in all acquired data (specialised and lipidic profiles with the same statistical importance – Figure 6B). However, this also highlighted that, even if in all OSMAC evaluated conditions a salt attrition effect was observed, however it is not constant and specifities could be observed in some culture media [69]. Thus, culturing the fungal strain on just one medium usually could not highlight a general effect of salinity [25, 70]. The most altered features corresponded to FAs (Table 3). Stahl & Klug reported that in fungi the most common and abundant FAs are 16:0, 18:0, 18:1, 18:2 that can represent up to 95% of the total FA content after analysing 100 strains of filamentous fungi including oomycetes, zygomycetes, ascomycetes and basidiomycetes [54]. Similar lipid main compositions were further confirmed in previously published data [31, 46, 55, 71-73]. However, lipids in fungi can be extremely diverse and more unusual compounds such as the polyunsatured FAs (PUFAs) 18:3, 20:4 or 22:6 can also been found to be produced in some conditions [46, 72, 74, 75].

Various studies have reported the impact of osmotic stress or salinity on membrane fluidity through modification of FAs saturation. In fact, the modification of the FA composition is essential for normal cell function in response to salinity change, mainly by maintaining fluidity or inducing rigidity of the cell membrane [76]. However, the observed effect remains different according to the fungal strains studied. In the literature, several studies in yeasts reported that an increase in salinity enhanced the rate of FA unsaturation (18:1, 18:2) [20, 62, 67, 77, 78]. In the case of *Debaryomyces hansenii* such an increase may appear only at a high level of NaCl [20]. In the other direction, such stress did not induce significant changes in the unsaturation of FAs as with the yeast *Yarrowia lipolytica* [15]. Contrarily, in filamentous fungi such as *P. notatum*, *P. chrysogenum* and *Aspergillus flavus* the presence of NaCl in culture media increased the abundance of saturated FAs (16:0, 18:0), while significantly decreasing the unsaturated ones [62]. One study focused on a marine derived *Epicoccum nigrum* also highlighted changes in the FA composition with the observation of a lower proportion of the unsaturated 18:1, 18:2 as an adaptive response to salinity stress in acidic conditions [77], even if this observation was strongly influenced by pH and temperature. In this way, our study was consistent with literature, where FA composition was clearly altered by the culture condition modifications evaluated [31, 46, 54, 73], where differences were observed not only in the diversity and types of FAs, but also in their relative concentrations. The most abundant FAs, whatever the culture conditions, were found to be 16:0, 18:0, 18:1 and 18:2, representing more than 90 % of the total FA content. The use of SSW did not modify the production of these four main FAs, but it increased the production of linear or branched minor even FAs, which, except the 20:5, all corresponded to saturated FAs (12:0, 14:0, *br*-16:0, *br*-18:0, 26:0). In comparison, FA features overproduced in DW conditions corresponded to the unsatured 19:1, 14:1 and *cj*-18:2, together with the saturated 17:0 DMA, 19:0 and 20:0. Altogether, this study further highlights the complex mechanisms involved in salinity adaptation in fungi.

In addition to FAs, two features were highlighted as significantly overproduced by the presence of SSW. They were putatively identified as dankasterone B and one of the isomeric cytochalasin B_5_, IV or 21,22-dihydrocytochalasin B. Dankasterone B is a steroid isolated from a marine-derived fungus *Penicillium* sp. which showed significant cytotoxic activity against HL-60, Hela, and K562 with IC_50_ values of 3.25, 4.74, and 7.89 µM, respectively [57]. Cytochalasin B_5_ and isomers are complex phenylalanine-derived macrolides belonging to the cytochalasan family, a class of fungal natural products produced by a large range of fungi from various ecological niches including marine. Cytochalasins all show various biological activities but are considered mainly as cytostatic or cytotoxic compounds that can be involved in chemical war against competitors [79]. However, most of the features for which a significant overproduction was observed under SSW conditions could not be annotated using the natural products databases queried. This result further demonstrates that salinity can induce specialised metabolism pathways and then the production of potential bioactive metabolites that can occur *in situ* but remain cryptic in artificial laboratory conditions [25, 80-84].

Regarding the metabolic consequences of using mussel flesh in the culture medium instead of yeast extract, a preliminary analysis of the data as simple as the unsupervised PCA clearly pointed out an effect of these environmentally relevant culture conditions. In addition, the use of the multiblock approach (Figure 5A) highlighted that each evaluated part of the metabolism was impacted by the OSMAC strategy. However, the most significantly altered part of the analysed metabolism was the FA composition of lipids. This result was unexpected as in a previous study focused on specialised metabolites it was shown that, when cultured in the MES-SSW medium, *P. restrictum* MMS417 overproduced two classes of specialised metabolites, namely pyran-2-ones and lactam macrolides [21]. It therefore appears that the most important response was firstly the induction of lipid metabolism pathways necessary for the physiological adaptation of the fungus, before the expression of more complex biosynthetic mechanisms of specialised metabolism.

As for salinity, mussel flesh had a minor effect on the production of the four major FAs 16:0, 18:0, 18:1 and 18:2. On the contrary, we observed drastic metabolic changes in the production of branched FAs (4,8,12-trimethyl-13:0, *iso*-17:0, *anteiso*-17:0, *iso*-16:0, *iso*-18:0, *anteiso*-18:0, dimethylacetal 18:0 and *br*-18:0) and long-chain FAs (20:1, 20:2, 20:4, 20:5, 22:1, 22:2) which are rather unusual in fungi [46, 75, 85-89]. Very interestingly, we observed under these conditions the presence of seven FAs which have never been described in fungi (4,8,12-trimethyl-13:0, *iso*-18:0, *anteiso*-18:0, dimethylacetal 18:0, *br*-18:0, 20:4, 20:5) but which have been previously reported to be produced by mussels [90, 91] (signals from blank mediums being removed in the data analysis process). This observation may be explained by (1) the presence of signal metabolites in the mussel extract that initiates such unusual FAs production, or more likely by (2) the presence of their precursors in the mussel extract that can be incorporated in fungal biosynthetic pipelines leading to complex lipids such as triglycerides or phospholipids that include these exogenous FAs. Such production of natural products through mixed biogenetic origins have been few demonstrated so far but it is hypothetised for the biosynthesis of brominated alkaloids from sponge and their endosymbiotic microorganisms [92], for the arginine-citrulline biosynthetic loop in the *Amphimedon queenslandica* holobiont [93] and for the sponge/bacterium mixed origin of calyculin A [94]. In our model, such phenomenon would be particularly interesting to confirm through targeted lipid purification or deep investigation of the two protagnonists genomes. It is also noteworthy that a feature annotated as agaric acid was significantly induced by the use of the MES medium. This tricarboxylic fatty acid, widespread in fungi [60, 61], is a competitive inhibitor of the adenine nucleotide translocase known to strongly inhibit FA synthesis [60, 95], alter sterol synthesis [60] and reduce bacterial biofilm formation [96]. Thus, it can be hypothesised that this compound could be involved in the regulation of the mussel bacterial population by *P. restrictum* within the holobiont microenvironment. In addition, in could be interesting to know if some other environmental stresses (other osmotic stresses [25, 97], microbial co-culture [27], etc.) ou the presence of other holobiont extract within the medium (oysters, sponges [98], seeweads [99], etc.) may induces similar responses on the lipid metabolism in comparison to mussel extract and/or salt presence.

In conclusion, this study engaged an ecologically-oriented OSMAC approach to expand fungal chemical diversity using not only specialised metabolite profiling but also lipid profiling and FA characterisation. All those data were explored, for the first time in this context, simultaneously using multiblock data mining strategies (COMDIM and MBPLS-DA). This approach was proven to be efficacious as it allowed to show that under some environmentally relevant conditions not only the specialised metabolism is strongly affected, but also the whole cellular physiology of the fungus as revealed by lipid alteration. Acquiring such a large quantity of chemical data on a single organism provides valuale insight to explore the biochemical basis of fungi in regards to interaction with the host after integration of complementary OMICS data (transcriptomic, proteomics, etc.) along with genome-scale metabolic modelling [100, 101].

This is in line with the observations made recently by Carroll *et al.* on the importance of using culture conditions close to the environment in which they naturally grow, when they observed that only 66% of the studies carried out on marine fungi in the last five years used cultures including seawater [102]. Host-derived media appear clearly more relevant than classical artificial culture media [21-24]. Such culture condition may not be considered as a strictly environmentally relevant. However, to the best of our knowledge, it remains currently one of the closest solutions to mimic host related condition at the laboratory level. In the present study, as we observed the presence of mussel FAs in the lipid composition of the fungus, we can assume that such lipids of hybrid origin are naturally produced within the bivalves, which could not have been observed otherwise. Furthermore, compounds rare or unknown in fungi were also observed, such as a number a FAs showing the lack of data on the fungal lipidome. Even more than for natural products, research into the chemistry of fungal lipids needs to be encouraged and continued.

Finally, it appears that the use of environmentally relevant conditions for the chemical study of marine microorganisms is essential not only for the discovery of new bioactive molecules [102], but also for understanding chemical interactions within marine microenvironments.

## Acknowledgments

This work is a part of the Ph.D. thesis of the first author Van Tuyen Le. Authors would like to thank the Corsaire-ThalassOMICS Metabolomics Core Facility of Biogenouest, Nantes Université for supporting LC-HRMS analyses.

## Author contributions

Olivier Grovel initiated the project. Samuel Bertrand designed the experimental works and carried out the multi-block analysis. Marion Brandolini-Bunlon performed the multi-block analysis validation. Aurélie Couzinet-Mossion and Gaëtane Wielgosz-Collin supervised the GC-MS experiments and their analysis. Vony Rabesaotra performed the GC-MS experiments. Van-Tuyen Le was involved in the collection of all data analysis and performed the all data analysis. Van-Tuyen Le, Samuel Bertrand and Emanuel Gentil perform the LC-HRMS analysis. Van-Tuyen Le and Thibaut Robiou du Pont performed fungal culture. Van-Tuyen Le and Samuel Bertrand wrote the draft manuscript. All authors revised, read, and approved the final version of the manuscript.

## Compliance with ethical standards

## Ethical approval

This article does not contain any study with human and/or animal participants performed by any of the authors.

## Supplementary data for

**Table S1.** Parameters used for automatic peak picking using MZmine 2.

| Step | Parameter | Positive ionization mode | Negative ionization mode |
| --- | --- | --- | --- |
| **1) Crop filter** |  |  |  |
|  | Retention time | 0.66-45 min | 0-45 min |
| **2) Mass detection** |  |  |  |
|  | Mass detection, mass detector | Centroid | Centroid |
|  | Noise level | 2.1E4 | 1.2E4 |
|  | Scans MS level | 1 | 1 |
| **3) Chromatogram builder** |  |  |  |
|  | Algorithm | ADAP Chromatogram builder | ADAP Chromatogram builder |
|  | Min group size in # of scans | 2 | 2 |
|  | Group intensity threshold | 6E4 | 2E4 |
|  | Min highest intensity | 2.1E4 | 1E4 |
|  | *m/z* tolerance | 30 ppm | 20 ppm |
| **4) Chromatogram deconvolution** |  |  |  |
|  | Algorithm | Wavelets ADAP | Wavelets ADAP |
|  | S/N threshold | 10 | 10 |
|  | S/N estimator | Intensity window SN | Intensity window SN |
|  | Minimum feature height | 20 | 100 |
|  | Coefficient/area threshold | 150 | 110 |
|  | Peak duration range | 0.0- 10 | 0.0-10 |
|  | RT wavelet range | 0.0 – 0.20 | 0.0 – 0.07 |
| **5) Isotope peak grouper** |  |  |  |
|  | *m/z* tolerance | 30 ppm | 20 ppm |
|  | Retention time tolerance | 1.8 min | 0.5 min |
|  | Maximum charge | 2 | 2 |
|  | Representative isotope | Lowest *m/z* | Lowest *m/z* |
| **6) Normalisation** |  |  |  |
|  | *m/z* tolerance | 30 ppm | 20 ppm |
|  | RT tolerance | 1.8 min | 0.5 min |
|  | Minimum standard intensity | 2E5 | 1E4 |
| **7) Join aligner** |  |  |  |
|  | Algorithm | Ransac aligner | Ransac aligner |
|  | *m/z* tolerance | 0.0 *m/z* or 30 ppm | 50 ppm |
|  | RT tolerance | 1.8 absolute (min) | 0.7 absolute (min) |
|  | RT tolerance after correction | 1.8 absolute (min) | 0.7 absolute (min) |
|  | Ransac iterations | 200000 | 20000 |
|  | Minimum number of points | 50.0% | 50.0% |
|  | Threshold value | 1 | 1 |
|  | Linear model |  |  |
| **8) Peak list rows filters** |  |  |  |
|  | Minimum peaks in a row | 4 | 4 |
| **9) Gap filling** |  |  |  |
|  | Algorithm | Peak finder (multithreaded) | Peak finder (multithreaded) |
|  | Intensity tolerance | 50% | 30 % |
|  | *m/z* tolerance | 30 ppm | 20 ppm |
|  | Retention time tolerance | 1.8 min absolute | 0.7 min absolute |

